# Reply To: Molecular Brightness analysis of GPCR oligomerization in the presence of spatial heterogeneity

**DOI:** 10.1101/822296

**Authors:** Michael R. Stoneman, Gabriel Biener, Valerică Raicu

## Abstract

Annibale and Lohse have recently suggested a way^1^ in which the two-dimensional fluorescence intensity fluctuation (2D FIF) spectrometry^2^ may be further refined. Their main suggestion is to include a step in the analysis process where a case-by-case inspection of individual regions of interest of a membrane allows for selection of portions of the membrane which are “as homogenous as possible” and thereby exclude intensity spots potentially related to other sub-cellular structures. By incorporating that proposal into an objective and reproducible algorithm, here we show that 2D FIF has a built-in capability to automatically filter out such contributions, and that further removal of inhomogeneities does not alter the final results.

---

We agree with the first observation made by Annibale and Lohse that the molecular brightness of a small region of a single cell expressing membrane-targeted monomeric eGFP (i.e. PM1-eGFP) is varying between nearby segments. To be meticulous, however, it must be stated that the range of brightness heterogeneity (i.e., a factor of 1.5) found for the small subset of data they analyzed falls within the brightness distribution assigned to monomers, as can be seen by assessing the positions of the black and red dotted lines in Fig. 1a relative to the entire distribution represented by the red dashed curve. That implies that for every brightness value higher than that corresponding to the center of the Gaussian (or its peak), there is a brightness value symmetrical about the peak, which obviously does not represent half of a monomer. Therefore, such a comparatively small variability in brightness should be at least in part ascribed to other causes than oligomerization, including variations in the micro-environment of the fluorescent probes (e.g., pH), orientation of the chromophores relative to the polarization of excitation light, and especially intrinsic membrane folds and creases (affecting the shape factor γ^2,3^), which can all lead to some broadening of the brightness distribution. That being said, larger fluctuations in the brightness (by factors of 2 or more) must indeed be present in the data in order for our brightness spectrograms to reveal larger oligomer sizes, as seen in Figure 1a herein as well as figures in our previous publication. We clarify further down below that such significant brightness distribution broadening is in fact facilitated by the inhomogeneous lateral organization of cell membranes, which causes local fluctuations in molecular concentrations, which are captured and exploited by FIF.

**Fig. 1.**
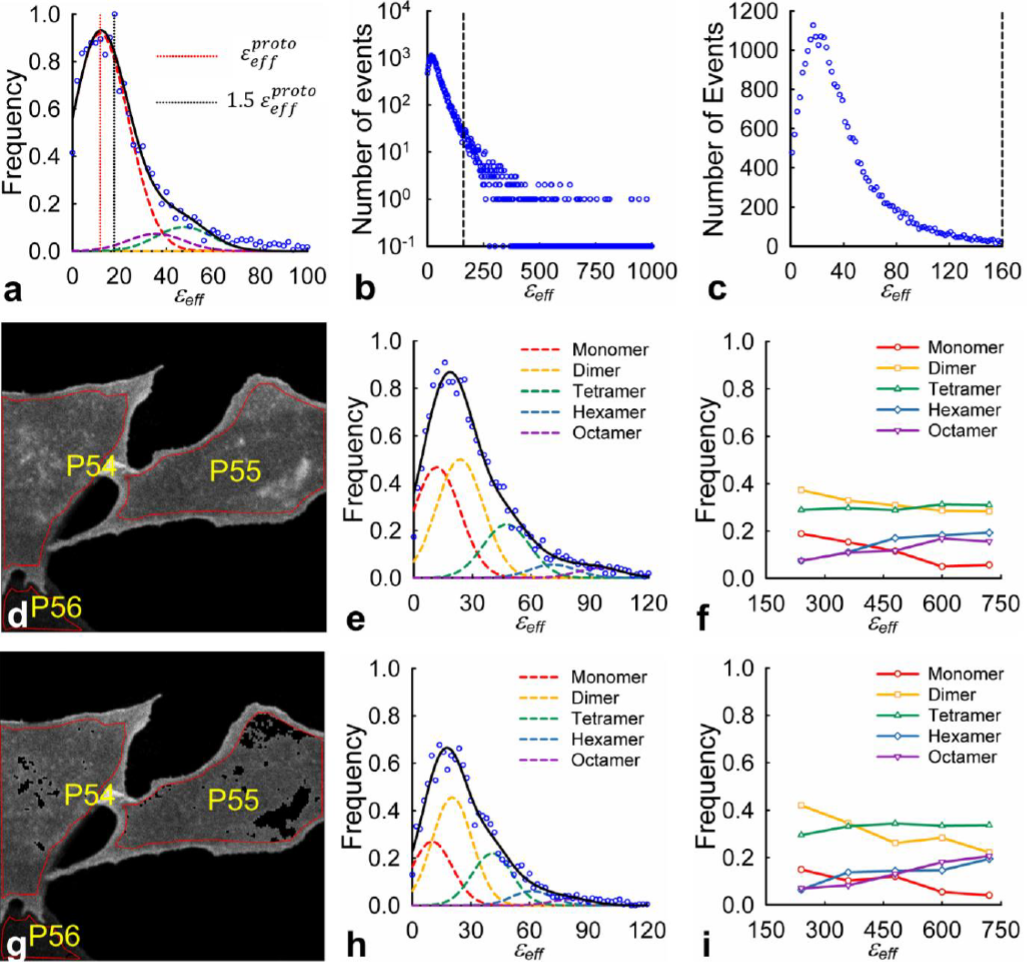
Effect of high intensity clusters on 2D FIF analysis. **a,** Effective brightness (*ε*_*eff*_) distribution obtained using FIF from cells expressing monomeric eGFP constructs (PM1-eGFP). **b.** Frequency distribution assembled from the complete set of brightness values obtained from images of CHO cells expressing wild-type secretin receptor treated with ligand published by Stoneman et al^2^. Brightness spectrograms were plotted for a range of *ε*_*eff*_ values between 0 and 160 (location indicated by the dashed black line). The data points to the right of the black dashed line originate from segments incorporating a cluster of high intensity pixels. **c,** Same distribution as shown in **(b)** with the y-axis plotted on a linear scale and cutoff at a brightness value of 160. **d-f**, 2D FIF analysis on fluorescence images of CHO cells expressing wild-type secretin receptor treated with secretin ligand. Regions of interest (ROI), indicated by red overlaid polygons in **(d)**, were segmented, and each segment analyzed to obtain a single *ε*_*eff*_ and concentration value. A brightness spectrogram constructed for a single concentration range (300-420 proto/μm^2^) is shown in **(e)**. The *ε*_*eff*_ distributions for each concentration were deconvoluted with a sum of Gaussians according to the 2D FIF approach to find the relative abundance of each oligomeric species, which was plotted as a function of concentration in **(f)**. The mean and standard deviation of the monomeric Gaussian used for the deconvolution was 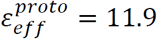 and σ=11.9, respectively (see Supplementary Fig. 1). **g-i**, 2D FIF analysis of the same dataset as used in row 2 applied after using the SLIC spot removal algorithm (see Methods) to remove clusters of high intensity pixels. The mean and standard deviation of the monomeric Gaussian used for the deconvolution was 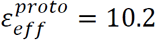 and σ=10.2, respectively (see Supplementary Fig. 1).

Regarding Annibale and Lohse’s suggestion to focus on regions of the membrane that are as homogenous as possible in order to avoid contaminating the data by intensity “hot spots” (associated perhaps with endocytic vesicles or other biological structures), we would like to emphasize that this is in fact an inherent feature of 2D FIF spectrometry, as already explained in our previous publication. In this method, dividing regions of interest into smaller segments leads to subsequent removal from brightness spectrograms of contributions from clusters with high average intensity, because the brightness of these regions is pushed to the far right of the spectrogram (see Fig. 1b). When fitting such spectrograms with Gaussian curves whose centers are *n* times the monomeric brightness, as it is done in 2D FIF, the amplitudes for Gaussians corresponding to very large *n* values are negligible (i.e., ~5/1000 or less). Hence, in 2D FIF the brightness spectrogram is cut off above that value (see Fig. 1c) in the process of data analysis. Therefore, 2D FIF automatically performs removal of high intensity spots using a rigorous low-pass filtering procedure, without the need for manual selection, which could be subjective.

We realize that the intrinsic filtering feature of 2D FIF may be a subtlety not immediately apparent to the casual reader, and therefore in this work we developed a computer program based on the simple linear iterative clustering algorithm (SLIC)^4^, to automatically remove high intensity clusters of pixels (see Supplementary Methods). The algorithm was first applied to cells expressing the monomeric construct PM1-eGFP (see Supplementary Fig. 1). Normalized frequency distributions were assembled from the brightness values obtained from the entire set of images both before and after applying the SLIC spot removal algorithm and were fitted to find the monomeric brightness value, 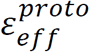, for both cases. Then, images of cells expressing the secretin receptor and treated with secretin ligand were analyzed using 2D FIF^2^ before and after application of the SLIC spot removal algorithm (see Fig. 1 d-i). As can be seen by comparing Fig. 1i to Fig. 1f, the hypothesis proposed by Annibale and Lohse that avoiding high intensity spots in the images will shift the interpretation of our data from secretin existing as a combination of various oligomer sizes towards that of a primarily monomeric species cannot be confirmed. The same analysis was performed on the set of images of cells expressing secretin receptor without any ligand treatment (Supplementary Fig. 2). There was no significant effect of the removal of the comparatively brighter spots from the analysis.

Parenthetically, the brightness values reported by Annibale and Lohse differ dramatically from our previously reported and current results – both in absolute values and as a trend –, perhaps due to possible errors in their data analysis procedure, as discussed in our Supplementary Note 1.

Nevertheless, the suggestion by Annibale and Lohse to focus on regions of the plasma membrane that are as homogenous as possible – which implies that there is a scale on which the distribution of proteins and phospholipids in the membrane is uniform – deserves further consideration. It is known that the membrane composition itself is inhomogeneous and presents lateral organization on all relevant size scales^5–8^. As one zooms in on a membrane there will always be another level of inhomogeneity, following a fractal or self-similar pattern^6^. Supplementary Fig. 3a provides additional support to this idea: As the image segment size shrunk, the brightness decreased, as expected for a bicontinuous (possibly fractal) distribution of proteins and phospholipids. This observation rules out Annibale and Lohse’s hypothesis that the membrane may be homogeneous on a certain small length scale, as far down as the length scale of the point spread function of the microscopes used (roughly 500 nm in diameter). It also offers an explanation as to why their simulations on homogenous mixtures of monomers and dimers could not be analyzed using the deconvolution procedure employed by FIF, as discussed further in Supplementary Note 2. While this is an interesting observation, which in itself could be a topic for larger publications, from the standpoint of a methods paper, a more relevant question is whether the measured brightness of the receptor of interest is independent of the length scale of the image segment. As seen in Supplementary Fig. 3b, the receptor oligomer size obtained from calibration against the monomeric construct (PM1) was highly reliable and scale-independent, provided of course that each data set was analyzed on the same scale. This finding is in full agreement with the 2D FIF results described above and in fact confirm the hypothesis that non-uniform lateral organization of the membrane coupled with concentration-dependent oligomeric size (according to the Law of Mass Action) allows FIF to detect large fluctuations in brightness and hence oligomeric size.

Finally, Annibale and Lohse propose that a combination of temporal brightness measurements, such as is done in the method of number and brightness analysis (N&B)^9^, with our method could provide additional information. Of course, we agree, in general, that a combination of the two techniques could be a powerful approach. However, we disagree with their assessment that all spatial distribution-based methods, including SpIDA and FIF, fail to discriminate large immobile background spots from large oligomers. SpIDA has been originally introduced to describe spatial intensity fluctuations by a pair of quantities consisting of single brightness value and a concentration of molecular entities (regardless of their oligomeric size)^10^. 2D FIF, by contrast, is a spectrometric method that provides entire probability distributions (i.e., spectrograms) of molecular brightness and concentration, not of entities, but rather of protomeric units that comprise them. As shown above, such spectrograms inherently filter out high intensity spots typically attributed to endocytic vesicles, clathrin-coated pits, etc., while allowing for fluctuations from segment to segment to provide sufficient statistics for determining the complete monomeric, dimeric, etc. contributions. This is the underlying working principle of FIF.

## Author contributions

M.R.S. performed data analysis and wrote the manuscript jointly with V.R and contributions from G.B. G.B. implemented the spot removal algorithms, performed data analysis, and contributed to writing of the methods section. V.R. designed and supervised the study, and wrote the manuscript jointly with M.R.S. and contributions from G.B.

## Competing interests

M.R.S., G.B., and V.R. have submitted a provisional patent application covering aspects related to the generation, analysis and applications of one and two-dimensional brightness spectrograms.

## Data Availability

Fluorescence images and ROI files used to generate the FIF spectrograms in this study have been deposited on the *Figshare* digital repository and are accessible from: https://figshare.com/s/77b90d060901fa8b4cb3

## Supplementary Methods: SLIC Spot Removal Algorithm

To remove intensity clusters in an image, we have turned to a computerized method that uses simple linear iterative clustering (SLIC)^1^ – which is used for ROI segmentation in the computer program provided by Stoneman et al^2^ – to generate segments of pixels which and then filters each segment based on average intensity. Initially, a region of interest (ROI) comprising the basal membrane of an individual cell is demarcated by the user. Then, the algorithm first breaks the ROI into smaller segment squares of length *l*_*s*_ pixels, and computes the center of mass, (*x*_*i*_, *y*_*i*_), of each segment *i*. Next, a pixel reassignment procedure is performed, whereby pixels are reassigned to a segment based on a distance criterion, *D*_*p*_, that takes its intensity, *I*, into account with an intensity weight factor, *W*_*I*_, along with the spatial distance between a given pixel, *p*, located at (*x*, *y*) and the nearest segment center (*x*_*i*_, *y*_*i*_). We set *l*_*s*_ to 7 pixels and *W*_*I*_ to 4096. The searching window was 14 pixels in length and max number of iterations was set to 5 (i.e., the loop between steps 3 and 9 will stop either when the criterion is obeyed or after 5 iterations).

A distribution of the average intensity of all the segments in a single ROI was then created and fit with a Gaussian function. Pixels within segments with average intensities greater than three standard deviations from the mean of the fitted Gaussian were set to 0. The filtered images were then segmented again, and each segment analyzed to obtain a brightness and concentration value, using the exact procedure and even chosen regions of interest from Stoneman et al^2^. The pixels which were set to 0 as a result of the filtering procedure were not included in the calculation of *ε*_*eff*_ and concentration.

The outline of the algorithm is provided below.

1. Segment ROI into squares of area 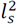
2. Calculate center of mass of each segment; remove segments with centers located outside ROI.
3. Initiate a label matrix, called *labeled_image*, where each pixel is ascribed an image segment index *i*.
4. **While** difference between centers positions in step k and step k-1 is greater than threshold, do
5. **For** each segment, *i*, centered at (*x*_*i*_, *y*_*i*_)
6. Calculate distances of each pixel within neighboring segments to segment centers (*x*_*i*_, *y*_*i*_) using (S28) from ref. 1.
7. If distance is smaller than the recorded one for the specific pixel, refresh distance value And mark the pixel as a member of the *i*^th^ image segment in the *labeled_image* matrix.
8. **End for**
9. Recalculate segments centers using all the pixels contained within a segment as marked in step 6.
10. **End While**
11. Calculate average intensity for all segments.
12. Assemble a histogram of average intensities.
13. Fit histogram with a Gaussian function.
14. Use the mean, *m*, and standard deviation, *σ*, of the fitting Gaussian to crop a new range between 0 and *m* + 2*σ*. Refit the new curve.
15. **If** ⟨*I*_*i*_⟩ > *m*_*new*_ + 3*σ*_*new*_
16. Set the pixels in the *i*^th^ segment to 0
17. Repeat entire algorithm for all ROIs.

⟨*I*_*i*_⟩ is the average intensity of the *i*^th^ segment while *m*_*new*_ and *σ*_*new*_ are the mean and standard deviation of the Gaussian fitting the new curve, respectively.

### Supplementary Note 1: Discrepancies in reported brightness and number values from Annibale and Lohse

In the Supplementary Table 1 of their correspondence^3^, Annibale and Lohse reported the results of their analysis of the intensity distributions extracted from four different ROIs (those shown in Fig. 1d of their correspondence) in three different ways: (i) They fit each of the four distributions with a Gaussian to extract the mean, ⟨*I*⟩, and standard deviation, *σ* values. These values are shown in columns 2 and 3 of our Supplementary Table 1. (ii) They used the original SpIDA algorithm^4^, without applying a correction for detector variance or intensity background, to extract a brightness value (*ε*_*corrected*_) for each of the four ROI. (iii) They used the SpIDA algorithm with a correction for detector variance (S=37) and intensity background value (*I*_*back*_ = 50) applied to again calculate brightness (*ε*_*corrected*_) for each of the ROI. We reanalyzed both our original data and their intermediate results and failed to reproduce the final brightness and number values, as discussed in more detail next.

There is a fairly straightforward relationship between uncorrected brightness values, *ε*_*uncorrected*_, calculated via the SpiDA algorithm, and the ⟨*I*⟩ and *σ*^2^ values extracted from the Gaussian fitting of the ROI intensity distributions, namely:

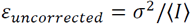

We have performed this simple calculation of *ε*_*uncorrected*_ for the mean and standard deviation values given for each of the four ROI and shown a side by side comparison of the values reported by Annibale and Lohse to those calculated with *σ*^2^/⟨*I*⟩ in Supplementary Table 1; as is evident from the table, the two sets of values are substantially different, with no observable correlation between the two.

We also calculated the brightness values from the mean and standard deviation of the four ROI after incorporating the correction for intensity background value and PMT noise variance. Adding in the correction for PMT noise should approximately amount to a constant offset value equal to the slope of the detector variance vs intensity plot (S), as given by the following:

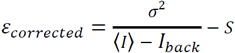

Shown in Supplementary Table 1 are *ε*_*corrected*_ for each of the four ROI in a side by side comparison to those reported by Annibale and Lohse, and again can find no correlation between the two sets of values.

We of course realize that these discrepancies do not really affect one of the main points made by Annibale and Lohse, that there is a heterogeneity in brightness values in closely spaced segments within a cell. As stated in the main text of this correspondence, we do not dispute this fact, which is why, in our original manuscript, the distribution of monomeric brightness values was represented with a rather broad Gaussian (mean 13.0 and standard deviation 13.4). However, the inconsistency in reporting brightness values does come into play when we address a second argument made by Annibale and Lohse. In Fig. 1f of their correspondence, they show that when decreasing the size of the ROI used to obtain intensity distributions, the calculated brightness values found from an ROI in a secretin-expressing cell becomes equal to the brightness value found from the monomeric EGFP sample. The implication of this finding is that secretin actually exists as a monomer, and artifacts due to inhomogeneities caused an overestimation of the secretin oligomer size in our original manuscript.

We have attempted to reproduce this effect using our entire set of data, with the results shown in Supplementary Fig. 3. Our analysis shows that no matter the size of the segment, there is a constant ratio (~1.25) between the average brightness value extracted from images of cells expressing secretin vs. the PM1-EGFP expressing cells. Two possibilities come to mind when trying to understand the origin of this erroneous effect that Annibale and Lohse proposed: (i) They used a small subset of image segments which showed this effect by pure statistical happenstance, but does not correctly characterize the entire dataset. (ii) There are errors being introduced to the brightness value calculations performed by Annibale and Lohse, which certainly seems possible based on our analysis of their Supplementary Table 1, which is presented in our Supplementary Table 1. It is also not clear where the values for *N* shown in Fig 1e as well as in SI Fig. 5 and of Annibale and Lohse come from, as they are an order of magnitude larger than the molecules/pixel reported in SI Table 1 of their correspondence, but an order of magnitude less than what would be expected for the total number of molecules in an ROI. This latter one is a less significant aside, but does add to the confusion regarding how the brightness calculations were performed by Annibale and Lohse.

### Supplementary Note 2: Simulated molecular brightness histograms lack complexity of membrane organization

In Supplementary Fig 5 of the correspondence by Annibale and Lohse, molecular brightness histograms are extracted from two different sets of simulated confocal microscopy images; the first set of images contains only a monomeric species, while the second set contains a 1:1 mixture of monomers and dimers. The resulting brightness histogram from the 1:1 mixture peaks at a brightness value of 9, which falls between that of the monomer distribution (~6) and what would be expected from a simulation corresponding to dimers (~12). Therefore, the brightness spectrogram of such a uniform mixture cannot be deconvoluted using a sum of the monomer and dimer brightness distributions. This result is not unexpected. The molecules in the simulations were distributed randomly, emulating a completely homogenous region in which a molecule could occupy each point in space with equal probability. However, in actual cell measurements, the distribution of phopholipids and proteins within the cell membrane itself is inhomogeneous, and most likely fractal and/or bicontinuous phase^5–7^. Because of this, local concentration fluctuations lead to fluctuations in oligomer size (via the Law of Mass Action) in each pixel. Since each image segment contains different portions of the bi-continuous distribution of phospholipids/proteins, the fluctuations in intensities are different from segment to segment, which is what we see in spectrograms from our original manuscript^2^. Furthermore, the stated concentration for each brightness spectrogram our original manuscript correspond in fact to a range of concentrations, which further broaden the spectrogram via changes in oligomer size, again governed by the Law of Mass Action.

## Supplementary Table and Figures

**Supplementary Table 1.** Comparison of recovered brightness values from four different ROIs to those reported in Supplementary Table 1 of Annibale and Lohse^3^.

| | $\langle I \rangle$ | $\sqrt{2}\sigma$ | $\epsilon_{uncorrected}$ | | $\epsilon_{corrected}$ | |
| --- | --- | --- | --- | --- | --- | --- |
| | | | Reported by<br>Annibale<br>and Lohse<sup>3</sup> | $\sigma^2/\langle I \rangle$ | Reported by<br>Annibale<br>and Lohse<sup>3</sup> | $\frac{\sigma^2}{\langle I \rangle - I_{back}} - S$ |
| Bottom left (3) | 1100 | 240 | 31 | 26 | 20 | -10 |
| Bottom right (4) | 1140 | 330 | 45 | 48 | 34 | 13 |
| Top left (1) | 1130 | 300 | 38 | 40 | 27 | 5 |
| Top right (2) | 1140 | 250 | 36 | 27 | 23 | -8 |

---

**Supplementary Fig. 1:**
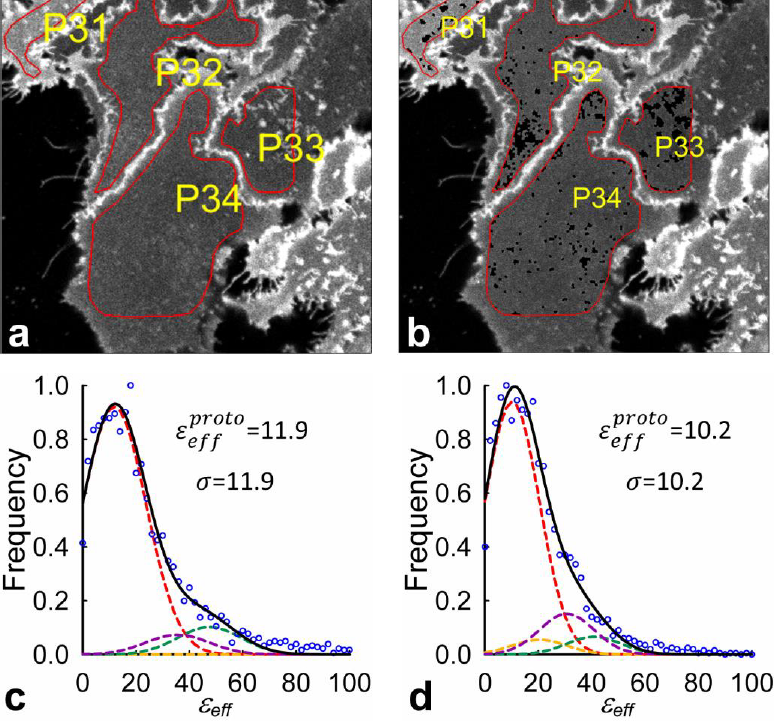
Illustration of the effect of removal of high intensity clusters from fluorescence images. **a,** Typical fluorescence image of Flp-In™ T-REx™ 293 cells expressing a plasma membrane-targeted mEGFP. Regions of interest (ROI), indicated by red overlaid polygons, were segmented, and each segment analyzed to obtain a single *ε*_*eff*_ and concentration value. **b,** Same image as in **(a)** after applying a spot removal procedure to remove clusters of pixels containing high intensities (relative to the intensities of neighboring regions within a cell). The spot removal procedure first used the SLIC method described in our original manuscript, to generate segments based not only on pixel location, but also intensity level. A distribution of the average intensity of all the segments in a single ROI was then created and fit with a Gaussian function. Pixels within segments with average intensity greater than three standard deviations from the mean of the fitted Gaussian were set to 0. The filtered images were then segmented again, and each segment analyzed to obtain a single *ε*_*eff*_ and concentration value. The pixels which were set to 0 as a result of the filtering procedure were not included in the calculation of *ε*_*eff*_ and concentration. **c,** The normalized frequency distribution assembled from the brightness values obtained from unfiltered images. The normalized distribution was fit with a sum (solid black curve) of Gaussians (dashed lines with various colors), to find the brightness of single 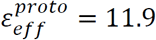. The various Gaussian peak positions were set to 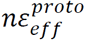, where *n* is the number of protomers in an oligomer, and their standard deviations, σ, were set equal to one another and determined from data fitting (σ=11.9). **d,** Same analysis as performed in **(c)** only to images filtered as described in **(b)**. The fitting resulted in a value of 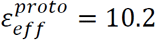 and σ=10.2.

**Supplementary Fig. 2.**
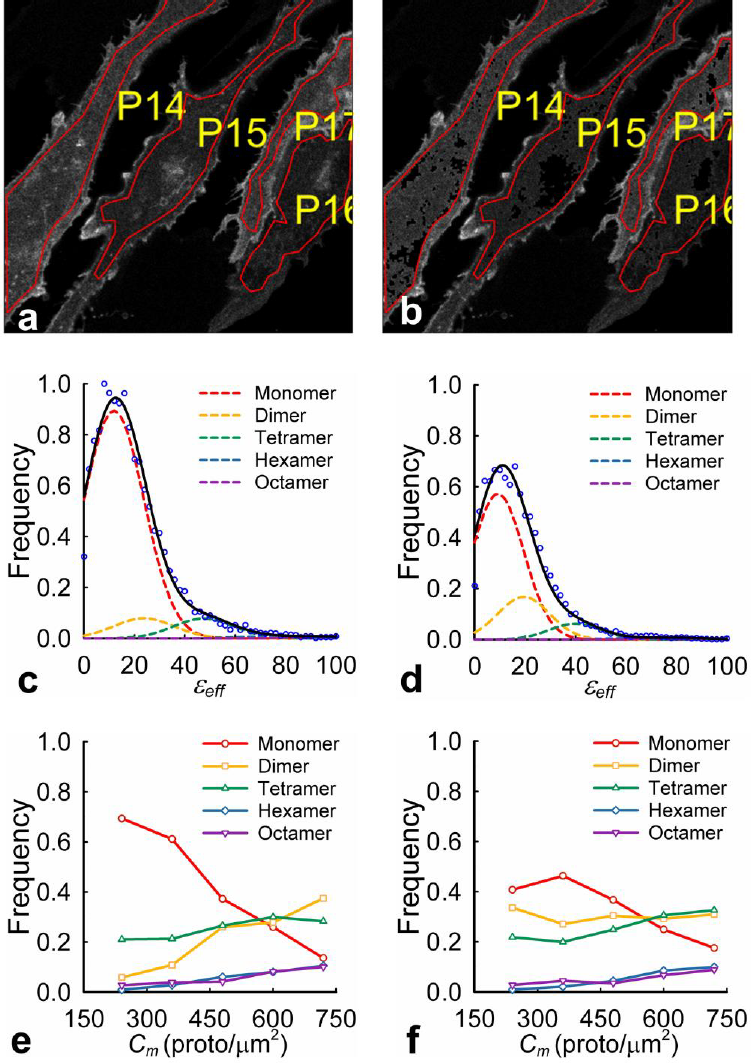
Investigating the effect of high intensity clusters on meta-analysis of brightness distributions. **a** Typical fluorescence image of CHO cells expressing wild-type secretin receptor (untreated). Regions of interest (ROI), indicated by red overlaid polygons, were segmented, and each segment analyzed to obtain a single *ε*_*eff*_ and concentration value. **b,** Same image as in **(a)** after applying a spot removal procedure to remove clusters of pixels containing high intensities, as described in the Supplementary Methods. **c, d** Brightness spectrograms constructed for a single concentration range (300-420 proto/μm^2^) for the unfiltered **(c)** as well as filtered **(d)** set of images. The *ε*_*eff*_ distributions for each concentration range was fitted with a sum of five Gaussians; the peak of each Gaussian was set to 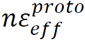, where *n* is the number of protomers in a given oligomer (e.g., 1, 2, 4, etc.). The value of 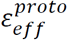 was found from analysis of the membrane bound monomeric eGFP data (see Supplementary Fig. 1 above); the values used were 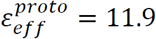 for the unfiltered images, and 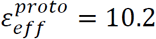 for those subjected to the spot removal procedure. Only the Gaussian amplitudes (*A*_*n*_) were adjusted in the process of data fitting which gave the fraction of protomers for each oligomeric species, i.e., *n*_*i*_*A*_*i*_/ ∑_*n*_ *nA*_*n*_. **e,f,** Relative concentration of prsotomers within individual oligomeric species vs. total protomer concentration, as derived by decomposing the spectrograms, like those shown in **(c)** and **(d)**, for each concentration range.

**Supplementary Fig. 3:**
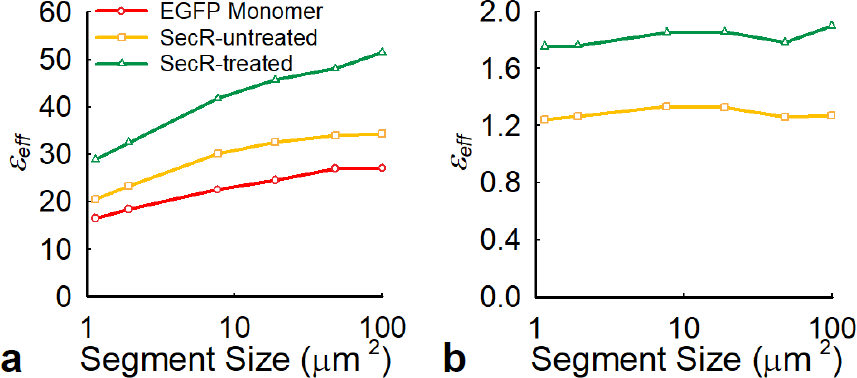
Average *ε*_*eff*_ of an entire distribution vs. segment size (i.e., analysis length scale). **a,** The average value of *ε*_*eff*_ was computed for brightness distributions prepared from three different datasets: CHO cells expressing wild-type secretin receptor (SecR) both (i) treated with ligand (green triangles) and (ii) untreated (yellow squares) as well as (iii) Flp-In™ T-REx™ 293 cells expressing a PM1-mEGFP (red circles). Each dataset was analyzed multiple times while varying the size of the segment used to obtain a single brightness/concentration pair as follows: 1.1 μm^2^ (289 pixels^2^), 1.9 μm^2^ (484 pixels^2^), 7.7 μm^2^ (1936 pixels^2^) 18.9 μm^2^ (4761 pixels^2^). The average *ε*_*eff*_ is computed for each of the distributions over all concentrations ranging from 0-1400 protomers/μm^2^ and *ε*_*eff*_ values ranging from 0-100, so as to avoid segments which produced extremely high *ε*_*eff*_, i.e. on the order of 10-100 times that of the monomeric brightness value. In our typical analysis, consisting of fitting the brightness spectrograms with a sum of Gaussians, these extremely high *ε*_*eff*_ values were automatically disregarded because they are significantly higher than the means of the Gaussians corresponding to the monomer, dimer, tetramer, etc. population. However, they do have a noticeable effect when using a simple averaging approach as we are doing in this figure. **b,** The average *ε*_*eff*_ values displayed in **(a)** were divided by the corresponding *ε*_*eff*_ value of the PM1-mEGFP sample obtained for that particular segment size. This shows that the receptor as well as calibration data from the monomeric sample are affected the same way and to the same extent by the length scale leaving unchanged the average size of the oligomers formed. The main body of the paper describes a more detailed analysis using the true 2D FIF method.

## Notes

https://figshare.com/s/77b90d060901fa8b4cb3

